# Enhanced maturation of human stem cell derived interneurons by mTOR activation

**DOI:** 10.1101/777714

**Authors:** Jianhua Chu, Megan L. Fitzgerald, Neha Sehgal, William Manley, Shane Fitzgerald, Harrison Naung, Ethan M. Goldberg, Stewart A. Anderson

## Abstract

The use of stem cell derived neurons for cell-based therapies is limited by a protracted maturation. We present a novel approach for accelerating the post-mitotic maturation of human stem cell derived interneurons via the activation of mTOR signaling. Lox sites were placed within *PTEN*, a key mTOR inhibitor, in a cortical interneuron (CIn) reporter line. Following directed differentiation and purification by FACS, the CIns were exposed to Cre-expressing lentivirus, then transplanted into mouse neocortex or plated onto cultured rat neocortex. Input synaptogenesis and dendritogenesis was greatly enhanced in the PTEN-deleted CIns. Whole-cell recording of the PTEN-deleted CIns in slices of transplanted neocortex revealed multiple indices of enhanced maturation. Finally, we observed similar effects using transient, doxycycline-inducible activation of AKT. We thus present an inducible, reversible approach for accelerating the maturation of human stem cell derived CIns, and to study the influences of this disease-related signaling system in human neurons.

## Introduction

The utility of human stem cell derived neuronal experimental systems is compromised by the protracted maturation time of human neurons. This challenge has been particularly problematic for inhibitory interneurons of the cerebral cortex (CIns). Pathology of CIns is implicated in major neurodevelopmental disorders, including autism, schizophrenia, and epilepsy (Inan et al., 2013; Jacob, 2016; Marin, 2012). Transplants of CIn progenitors from rodent embryos or mouse embryonic stem cells have demonstrated a remarkable capacity to disperse via migration and to differentiate into mature inhibitory neurons that modulate local activity (Southwell et al., 2014; Tyson and Anderson, 2014). These studies have also demonstrated the capacity of CIn transplants to affect symptom-related behaviors in rodent models of epilepsy, psychosis, chronic pain, and other disorders. However, CIn maturation extends well into postnatal life, continuing into the adolescent age range in both rodents and humans (Caballero et al., 2014; Wamsley and Fishell, 2017). Hence, despite the tremendous promise of this experimental system, there have been relatively few translational studies involving human PSC-derived CIns.

The maturation of human PSC-derived neurons has been accelerated through overexpression of progerin (Miller et al., 2013) and pharmacological inhibition of telomerase (Vera et al., 2016). Here, we use conditional and reversible activation of mTOR signaling to accelerate the maturation of CIns. mTOR is a broad modulator of proliferation, growth, and maturation (Saxton and Sabatini, 2017). In postmitotic neurons, activation of mTOR signaling enhances growth, connectivity, and action potential properties in a manner consistent with accelerated maturation (Maire et al., 2014; Vogt et al., 2015; Wyatt et al., 2014), but studies have traditionally interpreted these findings in the context of mTOR signaling-related neuropathology (Crino, 2016). Remarkably, we find that maturation of human CIns is accelerated in measures of morphology, input synaptogenesis, and electrophysiology by Cre-lox manipulation of the mTOR signaling repressor PTEN in post-mitotic interneurons. Importantly, transient activation of mTOR signaling via doxycycline-inducible expression of activated AKT yields similar results. We suggest that the capacity to controllably accelerate the maturation of human PSC-derived CIns will be broadly applicable in translational studies.

## Results

### A reporter line for the identification and isolation of newly postmitotic, migratory human CIn precursors

In both rodents and humans, major cortical interneuron subgroups originate from Nkx2.1-expressing progenitors in the subcortical telencephalon (Hu et al., 2017). As they exit the cell cycle, they begin to express Lhx6, and then maintain expression of this protein into adulthood (Lavdas et al., 1999). To generate an Lhx6 reporter line without grossly disrupting native Lhx6 expression, sequences encoding the fluorophore citrine were inserted into the Lhx6 gene, close to the native start site, followed by a 2A element (Fig. 1A-C). Differentiation of this line revealed citrine+ cells that co-label for Lhx6 immunofluorescence (Fig. 1D). FACS of these cells followed by co-culture with prenatal rat cortex showed differentiation into cells with morphologies of immature neurons that co-express nuclear Lhx6 (Fig. 1E).

**Figure 1.**
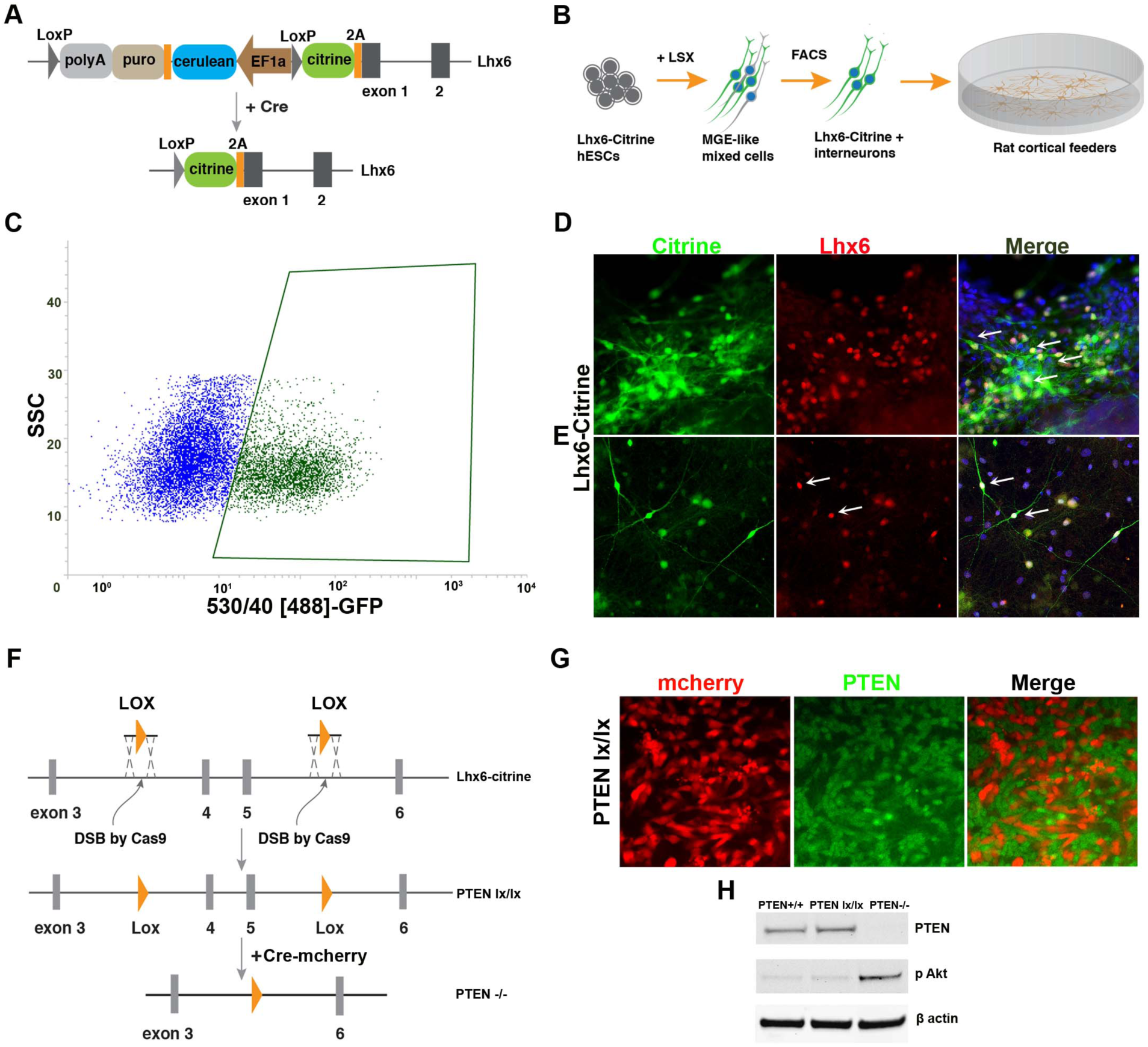
Generation of an Lhx6-citrine, PTEN^lx/lx^ hESC line. **A.** Strategy for generation of the Lhx6-citrine line. Citrine-2A is inserted at the N-terminus of Lhx6 using TALEN-based targeting of the initial coding region of Lhx6. A reporter line was generated by placement of citrine at the native start site for Lhx6 exon 1, followed by a p2A sequence and in frame read through into Lhx6 exon 1, which would not alter Lhx6 expression from this allele. **B.** Schematic of the differentiation to medial ganglionic eminence (MGE) mixed cells containing Lhx6-citrine+ precursors from day 0 (DD0) to DD35 according to already established protocol (Maroof et al., 2013). **C.** FACS of the Lhx6::citrine line differentiated till day 35. **D.** Representative immunostaining of interneuron precursors (depicted by white arrows) for citrine (green, using anti-GFP antibody) and Lhx6 (red, using an rabbit polyclonal antibody) at DD35 differentiated from from the Lhx6::citrine reporter hESCs. **E.** Representative immunofluorescent image of interneurons (depicted by white arrows) for citrine (green) and Lhx6 (red) grown on dissociated rat cortex cells for 6 weeks post-sorting. **F.** Schema of constructs used in generating PTEN lx/lx line from Lhx6-Citrine hESC reporter line by Cas9 and insertion of Lox sites. PTEN was knocked out from PTEN Lx/lx line by lentiviral introduction of Cre-recombinase. **G.** Representative immunofluorescent image of PTEN expression in cells from the PTEN lx/lx that were transfected with a mCherry-Cre-expressing lentivirus. **H.** Representative immunoblot of PTEN and pAKT of lysates of cells prepared from PTEN+/+, PTEN lx/lx, PTEN -/- lines. Data is represented for n = 5 independent experiments. Scale bars represent 20 μm.

### Loss of PTEN in post-mitotic interneurons accelerates maturation

The tumor suppression gene PTEN inhibits mTOR signaling and plays a pivotal role in regulation of neuronal growth, synaptogenesis, and the development of mature spiking characteristics (Kwon et al., 2006). To determine whether loss of function of PTEN accelerates the maturation of human stem cell derived interneurons, Lox sites were edited into introns 3 and 5 of PTEN (Fig. 1F) in an Lhx6-Citrine line, similar to the approach taken to generate PTEN^lx/lx^ mice. Introduction of Cre to these Lhx6Citrine:PTEN^lx/lx^ ESCs results in loss of PTEN immunofluorescence (Fig. 1G) as well as loss of PTEN protein and upregulation of phosphorylated AKT (Fig. 1H).

To study the effect of post-mitotic PTEN loss of function on hCIn development, Lhx6Citrine:PTEN^lx/lx^ ESCs were differentiated as per Fig. 1B with the following modifications: cells were replated after FACS, infected with an mCherry-Cre expressing lentivirus, then replated onto a two-week old carpet culture of E17.5 dissociated rat cortical cells. The sorted, transfected human Lhx6Citrine:PTEN^lx/lx^ ESCs were then grown for 6 weeks prior to fixation and immunocytochemical processing. Post-mitotic loss of PTEN resulted in a significant increase in dendritic complexity (Fig. 2A, quantified in Fig. 2B). Exposure to the mTOR signaling inhibitor rapamycin (50nM) reversed this increase (Fig. 2A, 2B). Like dendritic complexity, input synaptogenesis, defined by the counts of puncta of the excitatory postsynaptic marker PSD95 along mCherry+ dendrites in apposition to presynaptic terminals labeled with synapsin 1, was also greatly increased in PTEN^lx/lx^ Lhx6-Citrine+ cells (Fig. 2C, 2D). Again, rapamycin treatment reversed this, indicating that the effects of PTEN loss on dendritic complexity and input synaptogenesis are occurring largely through activation of mTOR signaling.

**Figure 2.**
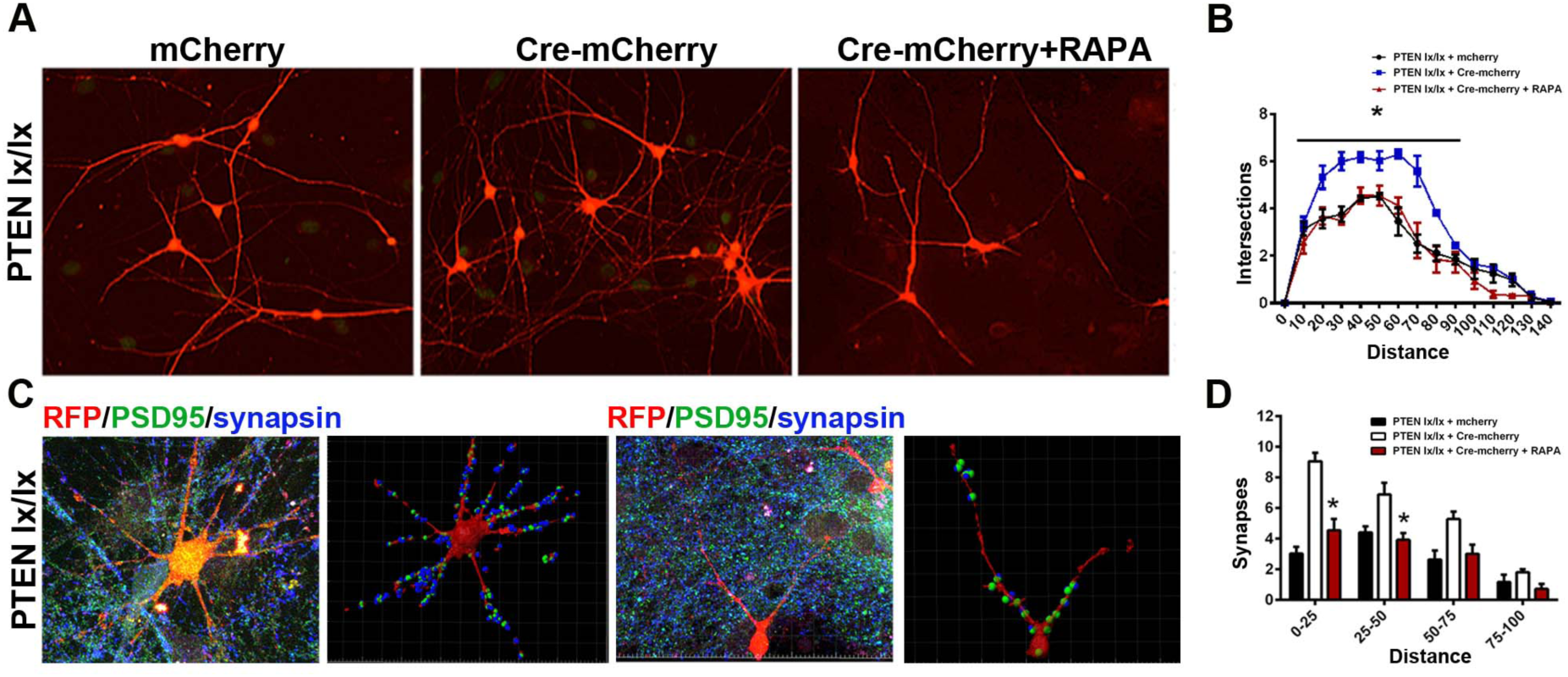
Loss of PTEN in post-mitotic cortical interneurons enhances morphological complexity and input excitatory synaptogenesis. Lhx6-citrine:PTEN lx/lx cells were differentiated as per Figure 1. FACS sorted cells were transfected with mCherry or mCherry-Cre lentivirus then grown on dissociated rat cortical cells for 6 weeks in the absence or presence of 50 nm rapamycin. **A.** Representative immunofluorescent images of interneurons from PTEN lx/lx stained with citrine (green) and PTEN (red). Relative to cells infected only with mCherry lentivirus (left panel), cells infected with Cre-mCherry (middle panel) appear substantially larger and have enhanced dendritic arborizations. The effect was inhibited by the mTOR inhibitor rapamycin (RAPA, right panel). **B.** Quantification of dendritic complexity by Sholl analysis. **C.** Left of each panel, collapsed z-stack confocal image showing mCherry (red), the excitatory postsynaptic marker PSD95 (green) and the presynaptic marker synapsin (blue). Right of each panel pair, reconstructions based on a 20-image stack using Imaris software to identify colocalizations of PSD95 with mCherry that include synapsin appositions. **D.** Quantification by IMARIS shows a significant increase in the number of synapses of PTEN lx/lx interneurons infected with mCherry-Cre lentivirus. The effect was reversed by rapamycin. Data is represented as mean ± SEM, n = 5 independent experiments; 20 neurons counted (*p < 0.05; ***p < 0.001). Scale bars represent 20 μm (A) and 40 μm (C).

While dendritic growth and complexity along with input synaptogenesis are hallmarks of neuronal maturation, electrophysiological characteristics define functionally mature neurons. To study these properties, Lhx6Citrine:PTEN^lx/lx^ ESCs were differentiated, subjected to FACS, replated and exposed to mCherry-Cre or mCherry control, and transplanted into the neocortex of neonatal immune-deficient (NSG) mice. After 6 weeks of maturation *in vivo*, cells were studied in cortical slices by whole-cell patch-clamp recordings (Fig. 3). PTEN KO cells had a resting membrane potential (*V*_m_) of −53.9 +/-10.7 mV; wild-type cells had a markedly more depolarized *V*_m_ of −30.6 +/-6.7 mV (Fig. 3C; unpaired t-test *p* < 0.001; *n* = 10 for both groups). In addition, PTEN KO cells had a membrane resistance (*R*_m_) of 644.1 +/-358.1 MΩ, whereas wild-type cells had an *R*_m_ of 3,602.1 +/-1,680.8 MΩ (Fig. 3C; *n* = 10 for both groups; *p* < 0.001). Action potential height was also significantly greater in the PTEN KO group.

**Figure 3.**
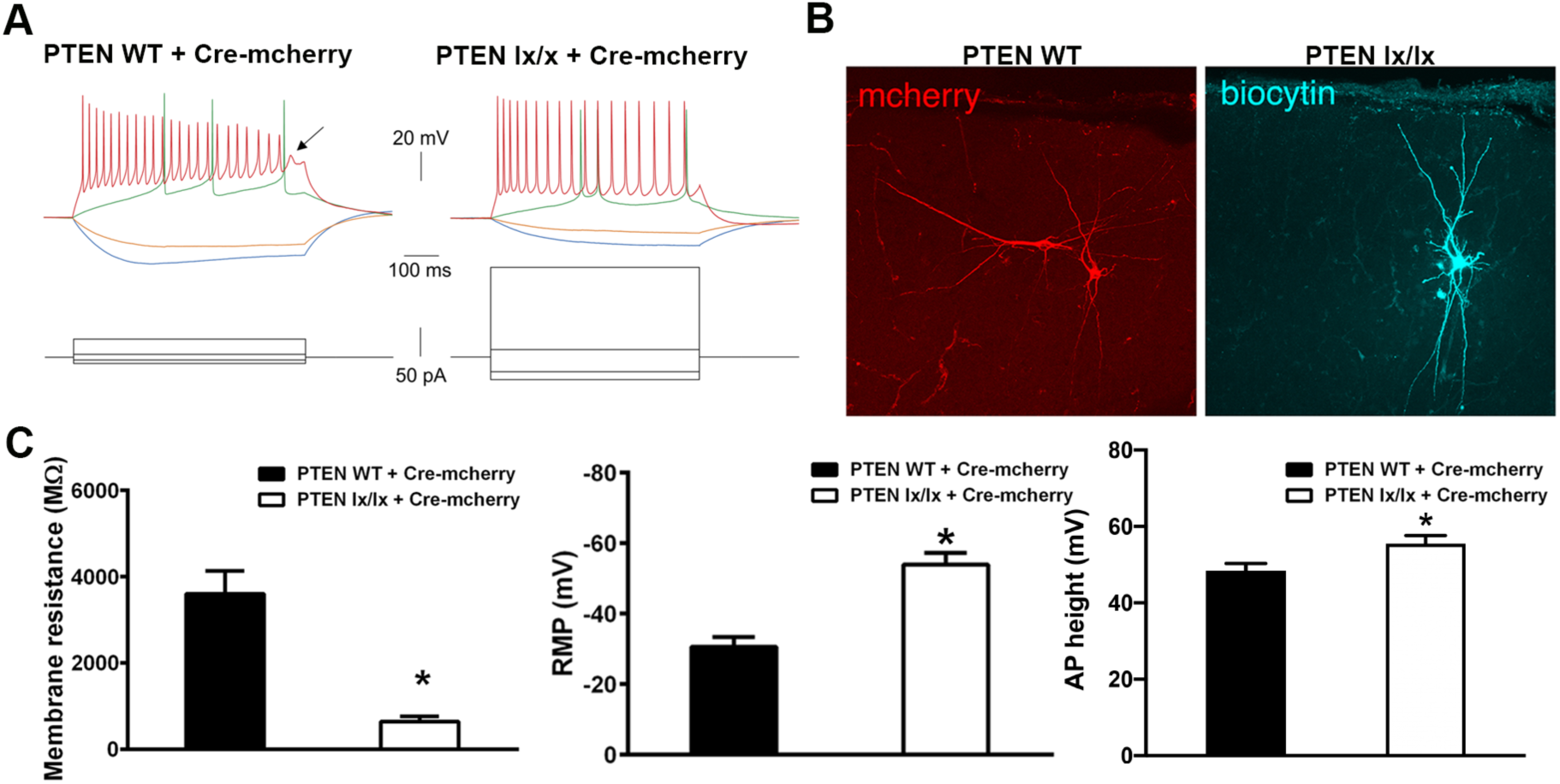
Loss of PTEN in post-mitotic hESC-derived interneurons enhances electrophysiological maturation. Cells were differentiated and infected as per Figure 2, PTEN WT line was generated from Lhx6-Citrine hESCs by CRISPR-Cas9 editing with no mutation introduced in the PTEN gene. Post-sorting citrine cells were infected with mCherry-Cre lentivirus followed by transplantation into the neonatal neocortex of NSG mice. Whole cell patch-clamp recordings were done 6 weeks post transplantation. **A.** Electrophysiological discharge patterns of PTEN WT and PTEN lx/lx stem cell-derived interneurons in culture. *At left*, firing pattern of a PTEN WT cell in response to −20 (*blue*), −10 (*orange*), +10 (*green*), and +50 pA (*red*) current injections. There was high input resistance (indicated by large hyperpolarization in response to small amplitude negative current injection) as well as spike height accommodation and failure (*arrow*) with sustained firing. *At right*, firing pattern of a PTEN lx/lx cell in response to - 40 (*blue*), −25 (*orange*), +15 (*green*), and +150 pA (*red*) current injections. There was markedly lower input resistance and more mature firing pattern with depolarizing current injection. **B.** Representative images of PTEN WT and PTEN^lx/lx^ cells, identified post-recording with immunofluorescence to mcherry and biocytin. **C.** The large differences in membrane resistance (left), and resting membrane potential (RMP), and the small but significant difference in action potential (AP) height were indicative of PTEN-/- cells being far more electrophysiologically mature. Data is represented as mean ± SEM, n = 10 cells per genotype.

Finally, we examined features of repetitive firing in response to square pulse depolarizations from −70 mV (using a hyperpolarizing DC current injection, so as to normalize across cells). Maximal steady-state firing frequency (SSFF) prior to AP failure during a 600 ms current injection was 26.5 ± 13.7 Hz for PTEN KO cells (*n* = 10) and 12.8 ± 17.0 Hz for wild-type (*n* = 8; *p* < 0.05). Maximal instantaneous firing frequency (IFF) was 64.5 ± 52.7 Hz for PTEN KO cells (*n* = 10) relative to 20.1 ± 15.7 Hz in wild-type (*n* = 8; *p* < 0.05). The values for SSFF and IFF for wild-type cells are artificially high because cells that did not fire, or did not fire more than one AP, were not included in the analysis. In sum, based on resting membrane potential, input resistance, action potential height, and repetitive firing, the loss of PTEN in post-mitotic hESC-derived interneurons results in a striking acceleration of electrophysiological maturation.

### Reversible activation of mTOR signaling accelerates neuronal maturation

The results presented above are strongly encouraging for the use of cell intrinsic, post-mitotic mTOR activation to accelerate the maturation of human stem cell derived interneurons. However, this approach irreversibly activates mTOR, a signaling pathway that broadly impacts cellular metabolomics, and could eventually produce deleterious phenotypes. We therefore sought an alternative, inducible/reversible approach for mTOR signaling activation.

Downstream of PTEN, AKT (protein kinase B) activates mTOR signaling. We acquired a doxycycline-inducible, constitutively active AKT and mCherry expression construct (TetO-myrAKT-mCherry-PGK-rTTA; Cellomics; Fig. S2). Lhx6-Citrine cells differentiated as per Fig. 1B were then subjected to FACS, infected with lentivirus containing the above construct, and replated onto prenatal rat cortical cells for 4 weeks (Fig. 4). Doxycycline (Dox, 100 ng/ml) was added to the cultures for 1, 2, 3 or 4 weeks (Figure 4A). The same Lhx6-Citrine line transfected with Cre-RFP was used as a control. Expression of doxycycline-dependent expression of mCherry and p-AKT was confirmed, with the highest levels produced by continuous addition of doxycycline as expected, but mCherry remained detectable following even one week of Dox addition (Fig S3). As with PTEN loss, activated AKT expression for four weeks resulted in tremendous enhancement of dendritic complexity and input synaptogenesis (Figs. 4B,4C,4D). Remarkably, while there was no effect of one week of Dox exposure, most of the benefit of activated AKT exposure for all four weeks was present when Dox was only added for the first two weeks. Taken cumulatively, these results suggest that transient activation of mTOR signaling via AKT can accelerate the maturation of human stem cell derived interneurons.

**Fig. 4.**
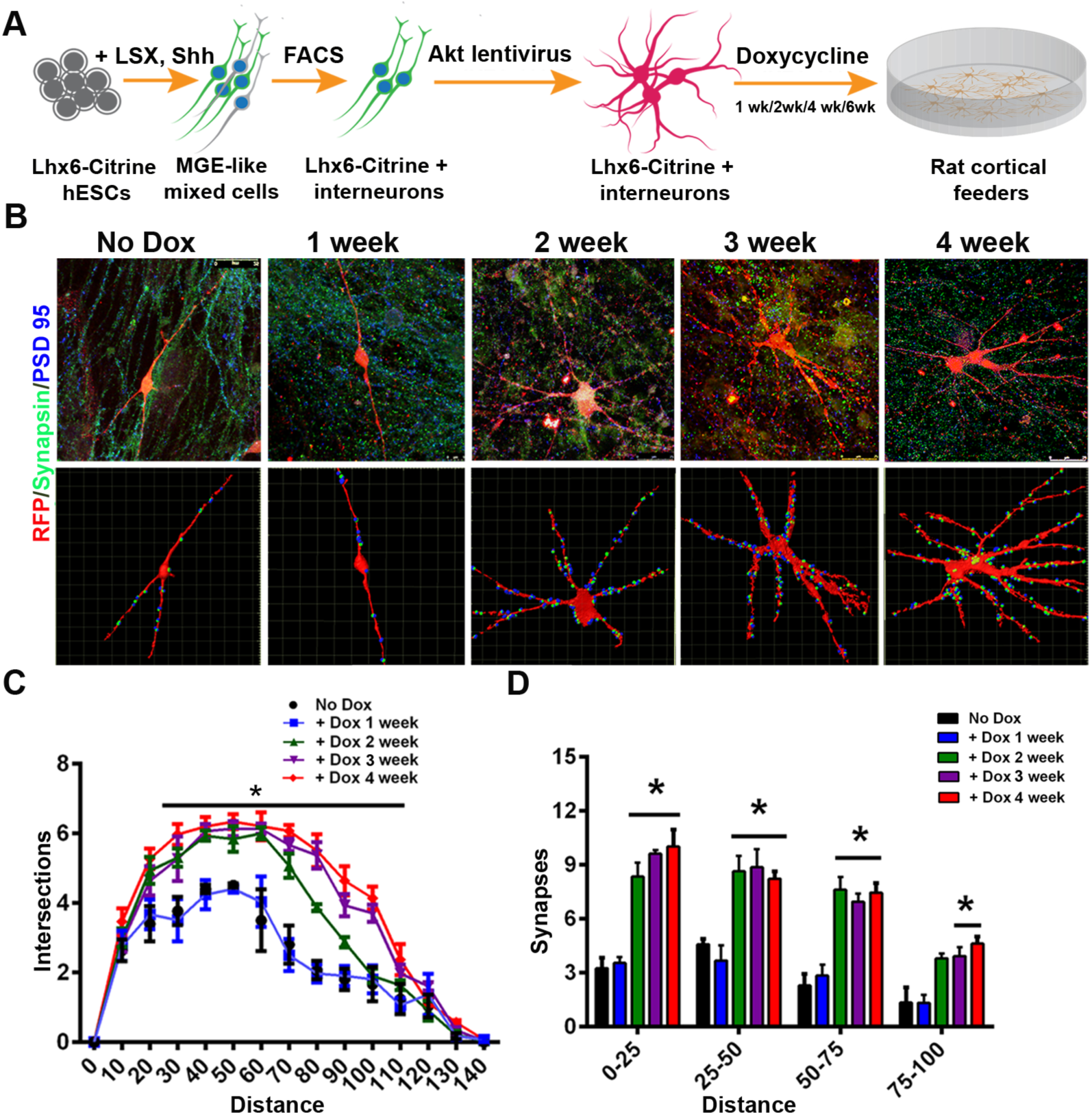
Transient activation of AKT signaling enhances interneuron maturation. (A) Schematic of the differentiation protocol, with and without the addition of doxycycline (Dox), for 6 weeks to the interneurons grown on cortical feeder cells. The steps from DD0 to DD35 were identical to those shown in Figure 1A. For these experiments, Lhx6::citrine cells were isolated by FACS on DD35, transfected with the mcherry::pLV-Tet-On-myr-AKT lentivirus and grown on rat cortical cells for 6 weeks. (B) Collapsed z-stack confocal image showing mCherry+ interneurons (red), DAPI (blue), the presynaptic marker synapsin (yellow) and the postsynaptic marker PSD95 (green) grown on rat cortical cells for 6 weeks after FACS analysis and transfection with mcherry::Cre lentivirus in the presence or absence of doxcycline for 1, 2, 3 or 4 weeks. (C) Quantification of neuron complexity by Sholl analysis. (D) Quantification of synapse density by IMARIS of interneurons after exposure to doxycycline. Data is represented as mean ± S.D, n = 5 independent experiments; 20 neurons counted (*p < 0.05; ***p < 0.001). Scale bars represent 40 μm.

## Discussion

The capacity to generate pluripotent stem cells from human somatic tissues, then to either induce or to direct the differentiation of those cells into disease-relevant neural subtypes, has engendered tremendous optimism that a tractable system for the study of human nervous system disease is at hand. While scientific advances based on hPSC-derived neurons and glia have indeed occurred, there is a major challenge to making actionable discoveries with this system: the naturally protracted timeframe of maturation of human neural cells. This challenge is especially relevant to the maturation of GABAergic CIns, a process that begins prenatally and extends into early adulthood (Anderson et al., 1995; Hu et al., 2017). Later defects in interneuron maturation and function are thought to contribute to some instances of schizophrenia and epilepsy (Inan et al., 2013).

This issue has become even more important with the revelation that the vast majority of neuropsychiatric disease-related single-nucleotide polymorphisms (SNPs) occur in non-coding regions (Avramopoulos, 2018). Due to chromosomal looping, the genes most linearly-proximal to a given variant are often not those directly affected (Mitchell et al., 2014). As these non-coding variants are generally poorly conserved across species, and the chromosomal looping evolves with cell type and developmental stage, it is ever more important to be able to study human neural cells at relatively mature stages of development (Mitchell et al., 2014; Norton et al., 2018).

To determine whether activation of mTOR signaling accelerates neuronal maturation, we first created an Lhx6-Citrine PTEN^lx/lx^ hESC reporter line which was then used to generate, identify, and isolate postmitotic human cortical interneurons. Loss of PTEN in these immature, postmitotic precursors resulted in increased excitatory synaptogenesis and morphological complexity, as well as electrophysiological measures of enhanced maturation. These effects were largely abrogated by addition of rapamycin, suggesting that mTOR signaling is primarily responsible for the maturational effects.

While these results represent an important proof of principle, irreversible activation of mTOR signaling is likely to produce deleterious cellular phenotypes due, for example, to suppression of mitophagy and autophagy (Baohan et al., 2016; Saxton and Sabatini, 2017). Although we have seen no evidence for reentry into the cell cycle or tumor formation, permanent/constitutive deletion of PTEN can also induce spiking abnormalities in neurons (Williams et al., 2015). We may be transiently engaging a similar mechanism to accelerate maturation of electrical excitability, with the main difference being that we have achieved exogenous control over PTEN expression via the dox-inducible genetic element. Altered PTEN and AKT signaling has been previously implicated in Alzheimer’s disease (Griffin et al., 2005) and autism spectrum disorders (Varga et al., 2009). To that end, the floxed PTEN cell lines developed in this study could prove useful for the study of disease-related mechanisms in post-mitotic cells.

To reversibly activate mTOR signaling, an inducible construct was used to activate AKT in newly postmitotic interneurons. Similar to loss of PTEN, AKT activation stimulated enhanced dendritic complexity and input synaptogenesis. Remarkably, even transient stimulation of mTOR signaling resulted in significant enhancement of dendritic complexity. Taken cumulatively, these results indicate that mTOR activation accelerates the maturation of newly post-mitotic neurons. Importantly, this is achieved in a cell autonomous manner such that maturational acceleration can be achieved after cells are transplanted in their immature, and in the case of CIns, highly migratory state. Transplanted PTEN^lx/lx^ hESC-derived interneurons exhibited profound acceleration of functional maturation as measured by standard indices including a markedly lower input resistance, more hyperpolarized (negative) resting membrane potential approaching mature CIns, and enhanced ability to discharge sustained trains of action potentials without spike height accommodation.

In sum, we have previously demonstrated that hPSC-derived CIns (hCIns) mature in their electrophysiological properties faster when cultured on dissociated rodent cortex than when either cultured on hPSC-derived cortical cells, or when transplanted into neonatal mouse brains (Maroof et al., 2013). We now find that hCIns growing on a carpet of embryonic rat cortical cells, then exposed to permanent or transient elevation of mTOR signaling, are far more mature than non-activated controls based their morphology, input synaptogenesis, and electrophysiology. Acceleration of maturation via inducible and reversible mTOR activation in other neuronal types could additionally facilitate their utility in cell-based therapies and in the study of later developmental, neurodegenerative, and senescence-related phenotypes.

## Acknowledgements

This research was funded by NIH NINDS K08 NS097633 and the Burroughs Wellcome Fund Career Award for Medical Scientists to EMG, and 2R01MH066912-16 to SAA.

## Author Contributions

JC, EG and SA designed experiments. JC, NS, MF, SF, BM, HN, and EG conducted experiments and organized data. JC, NS, MF, EG and SA wrote the manuscript.

## STAR Methods

### CONTACT FOR REAGENT AND RESOURCE SHARING

Further information and requests for resources and reagents should be directed to and will be fulfilled by the Lead Contact, Dr. Stewart Anderson

### EXPERIMENTAL MODEL AND SUBJECT DETAILS

#### TALENs and donor constructs

A pair of TALENs were engineered to recognize and target two 17-base pair sequences flanking the cutting site in exon 1 of the human *Lhx6* gene (left TALEN: tccccggcccatgtact, right TALEN: cgggcaacgccggggcg). The constructs were made by synthetic method at Genscript. The DNA of the insertion cassette (loxP-polyA-puro-cerulean-EF1a-loxP-Citrine-2A) was synthesized at Genscript and was cloned between two 1kb homology arms of the *Lhx6* gene.

#### Transfection and screening for targeted clones

Early passage H9 human ES cells (female) were used to generate the Lhx6-Citrine reporter line. 200-300K H9 ES single cells were plated on puromycin-resistant MEFs in one 6-well the day before transfection. 0.6ug donor plasmid and 0.2 ug each of TALEN plasmid were mixed with X-tremeGENE 9 (Roche) and added to the H9 ES cells.

Puromycin selection were performed 48 hrs after transfection at the concentration of 0.5ug/ml for 4-5 days. Puromycin-resistant ES cell colonies were hand picked and plated onto Matrigel-coated 96-well and cultured with mTeSR until confluent.

A pair of PCR primers (P9 and P13) located outside the two homology arms were used to amplify the targeted region. A predicted 5.7kb fragment corresponds to the targeted allele and 2.2kb the wild type untargeted allele. Several heterozygous clones and homozygous clones were identified from the screen and one homozygous clone (505-18) was used for further study. No detectable random integration of the donor plasmid was found in clone 505-18 by PCR. G-Band Karyotyping showed a normal 46, XX karyotype.

#### Characterization of Lhx6-Citrine reporter line

To validate the Lhx6-Citrine reporter line, 505-18 cells (after removal of the selection cassette by Cre mRNA transfection) was differentiated into mixed interneuron precursors according to (2). At dd35, the cells were co-stained with rabbit anti-Lhx6 and chicken anti-GFP (Abcam) antibodies for colocalization of LHX6 and Citrine proteins.

### Generation of Pten conditional ES cell line

#### CRISPR/Cas9 Plasmid

Several 20-nt target sequences were identified in intron 3 and 5 of the Pten gene. Each sgRNA sequence was functionally tested using Surveyor assay. Two of the sgRNA oligos (Pten-sgRNA-L4 and Pten-sgRNA-R1) were cloned between the two BbsI sites of pSpCas9(BB)-2A-GFP (PX458) plasmid according to (Ran et al., 2013).

#### Donors

Single-stranded oligodeoxynucleotides donors consists of two 60-nt homology arms flanking a 34-nt lox sequence were synthesized at IDTDNA.

#### Transfection and screening for correctly targeted clones

Two rounds of gene targeting were performed to insert two lox sequence (L-lox and R-lox) into the intron 3 and 5 of Pten gene respectively. For the first round, Lhx6-Citrine ES cells were transfected with pSpCas9-(Pten-sgRNA-R1)-2A-GFP and Pten R-1 ssODN using Nucleofector 2b device. After transfection, cells were plated onto 24-well Matrigel-coated plates in the presence of Y-27632. GFP-positive cells were isolated 24 hours post-transfection with FACS and plated onto MEF feeder plate. ES cell colonies that emerged after about 7 days were hand-picked and plated onto Matrigel-coated 96-well and cultured with mTeSR until confluent.

To screen for the correctly targeted clones, a pair of PCR primers (Pten-P5 and Pten-P2) were used to amplify the targeted region in intron 5 of Pten. A predicted 407-bp fragment corresponding to the correctly targeted allele were identified in several clones and one of the homozygous clones (505-18-12) was verified by sequencing and was used for 2nd round targeting.

The second round of targeting was performed similarly. 505-18-12 cells were transfected with pSpCas9-(Pten-sgRNA-L4)-2A-GFP and Pten L4 ssODN and correctly targeted clones were identified by PCR using Pten-P7F and Pten-P40R. A homozygous clone (505-18-12-2) was verified by sequencing for the insertion of lox sequence in intron 3 of Pten.

#### Characterization of Pten-flox/flox ES cell line

To further validate the established Pten-flox/flox line, 505-18-12-2 (Pten-flox/flox) ES cells were transfected with Cre mRNA to remove the ∼7.3 kb fragment flanked by the lox sites, which contains exon4 and 5 of Pten. The resulting Pten-/-ES cells were tested by Western blotting for the expression of PTEN protein using rabbit anti-PTEN antibody (Cell Signaling Technology). No detectable PTEN protein was found in the Pten-/-ES cells (Fig 2b).

A Ubc-mCherry-IRES-Cre lentivirus was used to infect Pten-flox/flox ES cell culture. ES cell colonies were fixed and stained for mCherry and PTEN expression 48 hours after infection. PTEN was found to be eliminated in mCherry positive cells but not in mCherry negative cells (Fig 2c), indicating the effectiveness of the Cre lentivirus as well as validating the Pten-flox/flox cells.

### METHOD DETAILS

#### Differentiation of cortical interneuron precursors

Human embryonic stem cells (hESCs) (H9) were grown on were plates coated with Matrigel (Corning) in StemMACS iPS-Brew XF (Miltenyi Biotec) supplemented with 10 μM Y27632 (Stemgent). Cells were passaged using Accutase (Sigma-Aldrich). Neural induction of the H9 cells occurred from differentiation day 0 until day 10. Cells were treated with “LSX” media consisting of 100 nM LDN-193189 (Stemgent), 10 uM SB431542 (Stemgent), and 2 uM XAV939 (Stemgent) in DMEM:F-12 (Thermo Fisher) supplemented with 20% Knockout Replacement Serum KSR (Invitrogen), 1% MEM non-essential amino acids (Invitrogen), 0.5 mL of 55 mM beta-mercaptoethanol, and 1X Glutamax (Gibco). LSX media was changed daily from day 1 to 4. On day 5 media was changed to a mixture of 3:1 “LSX”:”N2” media. “N2” media consisted of DMEM:F-12 (Thermo Fisher) supplemented with 0.75g of D-glucose (EM Science), 1% N2 supplement (Stemgent), 0.5 mL of 55 mM beta-mercaptoethanol, and 0.2% Primocin (Invivogen). “N2” stock media was further supplemented with B27 (without vitamin A) media (1:50)(Gibco), 1 μM purmorphamine (EMD Millipore), 10 μM Y27632 (Stemgent) and 100 ng/ mL human recombinant Sonic Hedgehog (R & D). On day 7, media was changed to 1:1 “LSX”:”N2” media. On day 9, media was changed to 1:3 “LSX”:”N2” media. From day 11 until day 18 cells were treated with “N2” media exchanged every other day. Starting at day 20 cells were treated with “NBM” media from day 18 to day 35. “NBM” media consisted of neurobasal medium (Gibco) supplemented with 1% Glutamax (Gibco), 0.75g of D-glucose (EM Science), and 0.2% Primocin (Invivogen). “NBM” stock media was further supplemented with 2% B27 with vitamin A (Gibco). Media was exchanged every other day.

#### Transplantation

Citrine positive Pten +/+ or Pten lx/lx cells were purified by FACS and plated onto Matrigel-coated 4-well plates in NBM/B27 media at 2 million cells per well. After 24 hours, 2ul Ubc-Cre-mCherry lentivirus (10^9 TU/ml) was added to the well in the presence of 5ug/ml polybrene. 48 hours post infection, the cells were collected and resuspended in NBM/B27 media at the concentration of 30K/ul. 1.5 ul of resuspended cells were transplanted bilaterally into the cortical plate of the somatosensory neocortex of P1 or P0 NSG neonatal mice at the following coordinates from bregma: 2.0 mm anterior, 1.0mm lateral and 0.5 mm deep.

#### Slice Preparation

Acute brain slices were prepared from mice (8-12 weeks) using standard techniques essentially as previously described (Goldberg et al., 2011). Mice were anesthetized via inhaled isoflurane and decapitated. The brain was rapidly removed to oxygenated, ice-cold artificial cerebrospinal fluid (ACSF) that contained, in mM: 87 NaCl, 75 sucrose, 2.5 KCl, 1.25 NaH_2_PO_4_, 26 NaHCO_3_, 10 glucose, 0.5 CaCl_2_, 4 MgSO_4_. Slices (275 μM thick) were cut on a Leica VT1200S and incubated in cutting solution a holding chamber at 32° C for approximately 30 minutes followed by continued incubation at room temperature prior to electrophysiological recording, at which point slices were transferred to a submersion type recording chamber attached to the microscope stage. ACSF used for recording contained, in mM: 125 NaCl, 2.5 KCl, 1.25 NaH_2_PO_4_, 26 NaHCO_3_, 10 glucose, 2 CaCl_2_, and 1 MgSO_4_. The solution was continuously bubbled with 95% O2 and 5% CO2 throughout cutting, slice incubation, and recording, so as to maintain a pH of approximately 7.40.

#### Electrophysiological recordings

Transplanted cells were identified via GFP expression, subsequently visualized using a 40X, 0.8 NA water-immersion objective (Olympus) on an Olympus BX-61 upright microscope equipped with infrared differential interference contrast optics. Current clamp recordings of transplanted cells were performed using the whole-cell patch clamp technique with electrodes pulled from borosilicate glass to a tip resistance of 5-7 MΩ using a Sutter P-97 pipette puller. Access resistance (R_s_) was < 25 MΩ upon break-in; data obtained from a given cell was rejected if Rs changed by > 20% during the course of the experiment. Internal solution contained, in mM: potassium gluconate, 130; potassium chloride, 6.3; EGTA, 0.5; MgCl_2_, 1.0; HEPES, 10; Mg-ATP, 4; Na-GTP, 0.3; biocytin, 0.1%. Osmolarity was adjusted to 285-290 mOsm using 30% sucrose. Voltage was recorded using a MultiClamp 700B amplifier (Molecular Devices, Union City, CA), lowpass filtered at 10 kHz, digitized at 16-bit resolution (Digidata 1440) and sampled at 50 kHz. pCLAMP 10 software (Axon Instuments) was used for data acquisition, and analysis was performed using the Clampfit module of pCLAMP and custom Matlab scripts. Resting membrane potential (*V*_m_) was defined as the mean membrane potential during a 10-second baseline current clamp recording; membrane resistance (*R*_m_) was calculated as the slope of the linear fit of the voltage/current relation near *V*_m_; action potential half-width (AP ½-width was calculated as the width of the AP at half maximal amplitude from AP threshold to peak of the AP; spike frequency adaptation was calculated as the ratio of the 1st to the 10th (*ISI*_1_/*ISI*_10_) or last (*ISI*_1_/*ISI*_n_) peak-to-peak interspike interval.

### QUANTIFICATION AND STATISTICAL ANALYSIS

#### Sholl and synaptogenesis analyses

Imaris (Bitplane) imaging and reconstruction software was used to create 3-dimensional representations of mCherry+ neurons and synaptic inputs. Each neuron was manually traced and automatically thresholded and masked. Artifacts were manually traced and excised. For Sholl analysis, the ‘Filament’ function was used to manually trace neuronal processes. Traced neurons were then analyzed using the ‘Filament No. Sholl Intersections’ function, and data was exported for further analysis.

To identify points of synaptic contact, the ‘spot creation’ function with an estimated XY diameter of 0.6 μm was used to label puncta of synaptic staining on each channel for PSD-95 and synaptophysin. ‘Spots’ were colocalized using the ‘colocalize spots’ function in Imaris. Threshold value was set to 1. The number of ‘spots’ on the presynaptic channel (synaptophysin) were recorded and used in the downstream statistical analysis where they were used to represent synapses onto the target neuron.

#### pAKT and mCherry quantification

To quantify immunofluorescence, the MBF Stereo Investigator system was used. The Optical Fractionator workflow probe was used to sample regions of interest (ROIs). Cells were chosen randomly prior to imaging. Once ROIs were traced and cells were chosen, images were captured at the same exposure time at 20X magnification. Cell fluorescence was then measured using Stereo Investigator’s pixel brightness analysis measurement. Each cells’ brightness was then averaged to determine average pAKT or mCherry expression for each ROI.

#### Statistical analysis

GraphPad Prism was used to conduct statistical analyses. Two-way ANOVAs were used to compare Sholl analysis and input synaptogenesis data between PTEN-/- and control lines, and between doxycycline-inducible AKT construct and control lines. Outliers were excluded. Unpaired T-tests were used to compare electrophysiology data between groups. Significance was determined at p < 0.05.

## DATA AND CODE AVAILABILITY

Dataset can be accessed through Mendeley: Fitzgerald, Megan (2019), “Enhanced maturation of human stem cell derived interneurons by mTOR activation”, Mendeley Data, V1, doi: 10.17632/jw9n4k9gny.1

No new code has been generated in this paper.

