## Supplemental for "Enhanced maturation of human stem cell derived interneurons by mTOR activation"

**Supplemental Table 1:**

Summary of electrophysiological characteristics

| | | Intrinsic properties | | Properties of individual action potentials | | Repetitive action potential discharge<br>Maximal firing frequency | |
|---|---|---|---|---|---|---|---|
| | ***n*** | ***V<sub>m</sub>*** (mV) *** | ***Rm*** (Mohms) *** | **Height** (mV) * | **½ width** (ms) | **Hz** *Steady-state* * | **Hz** *Instantaneous* * |
| *PTEN-WT* + Cre-mCherry | 10 | -30.6 ± 10.7 | 3602 ± 1681 | 48.4 ± 5.9 | 2.8 ± 1.0 | 12.8 ± 17.0 | 20.1 ± 15.7 |
| *PTEN*-flox/flox + Cre-mCherry | 10 | -53.9 ± 10.7 | 644 ± 358 | 55.2 ± 6.7 | 2.0 ± 0.7 | 26.5 ± 13.7 | 64.5 ± 52.7 |

**, p<0.05; **, p<0.01; ***, p<0.001*

**Supplemental Table 2:**

| **Oligonucleotides** | **Sequence** |
|---|---|
| Lhx6-P9 | aggctgtcatggtgtctgtagg |
| Lhx6-P13 | cgtggctcttgaagtaatcgc |
| Pten-P5 | acagcagtatcaggctgtaga |
| Pten-P2 | ccttactgccagacaacacatc |
| Pten-P7F | ggtggaaggagggcatttat |
| Pten-P40R | gccagcattcctacaagagc |
| Pten-sgRNA-R1 | tctcttcatcagtatgcgct |
| Pten-sgRNA-L4 | gaatgggggttggtaggata |
| Pten R1 ssODN | actggtaaagacagaagctctccatgtcatggtcacccactg ctctcttcatcagtatgc ataacttcgtata gcatacat tatacgaagttat gctagggtgcaagactgtcatcaatcctgaaagaaatcttatc tcttgttgtgaaagtat |
| Pten L4 ssODN | Gggagcaaatatgctgggactagtaatgataaatatgctcag ggaatgggggttggtagg ataacttcgtata gcatacat tatacgaagttat Ataaggatgaagactttcttcttaaattaaaatctctgataaatg ccctccttccaccta |

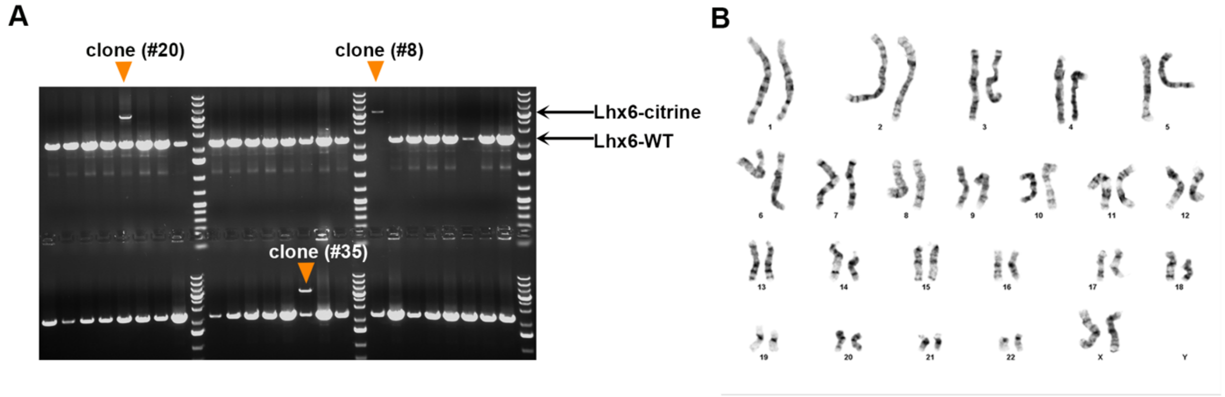

**Supplemental Fig. 1. Characterization of Lhx6-Citrine hESCs line.**
(A) PCR-based screening results of 48 hESC clones. A 5.7kb and 2.2kb products (black arrow heads) correspond to the targeted allele and the unedited allele respectively. Representative homozygous clone (#8) and 2 heterozygous clones (#20 and 35) are shown. (B) Normal 46 XX karyotype of line 505-18 by G banding.

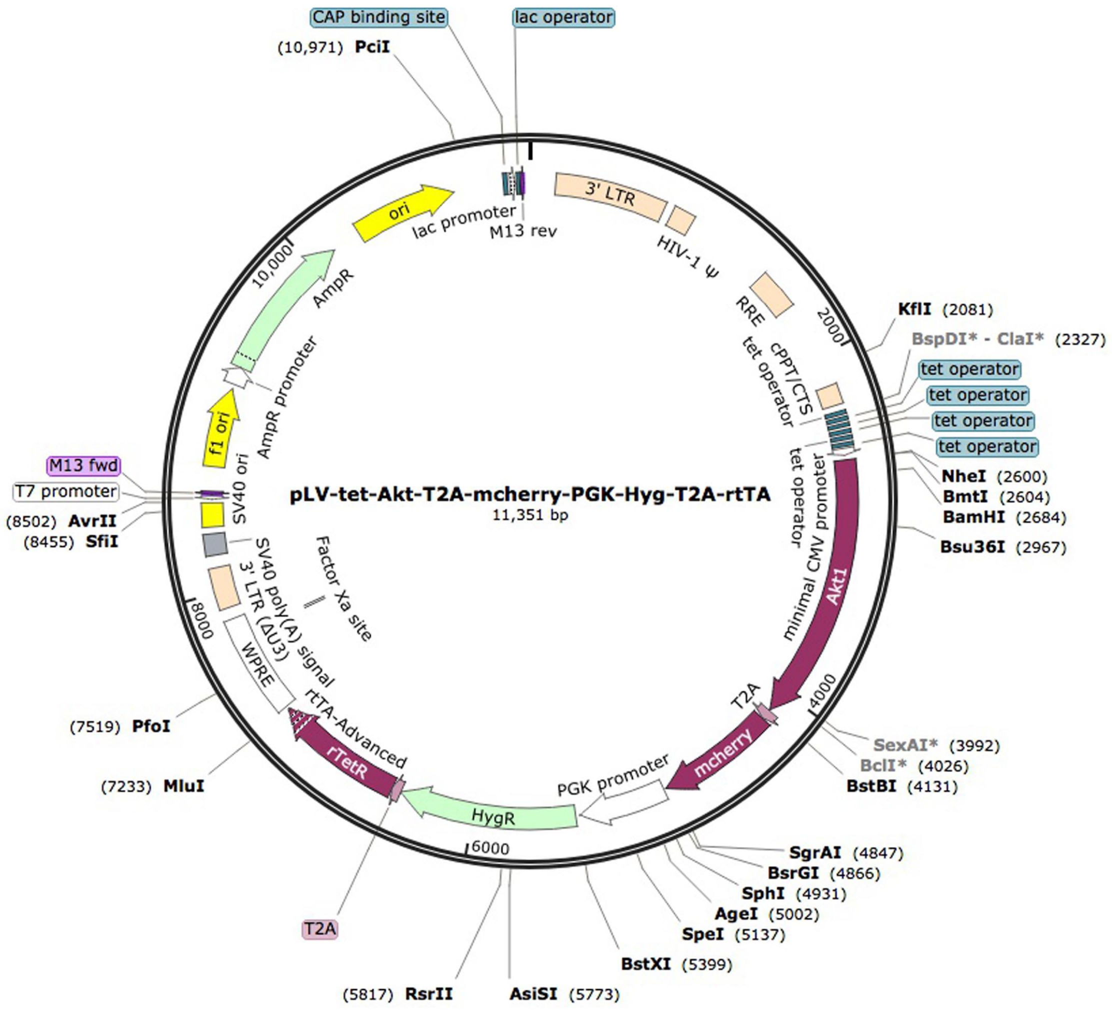

**Supplemental Figure 2.** A cDNA containing a src myristolation sequence and a human AKT1 coding region was cloned between the NheI and BstBI site of pLV-tet-T2A-mcherry-PGK-Hyg-T2A-rtTA vector to allow doxycycline dependent expression of constitutively active AKT.

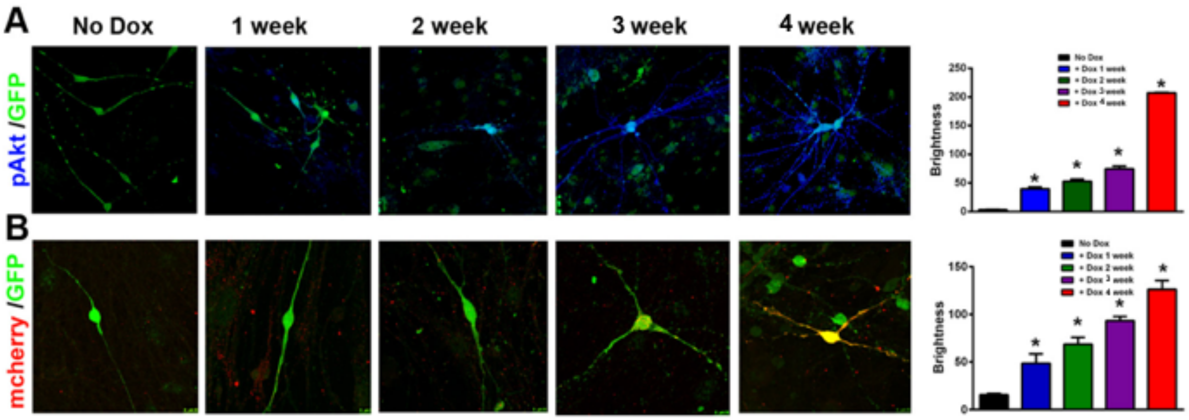

**Supplemental Figure 3. Expression of p-AKT and cherry after doxycycline exposure**. Lhx6-Citrine + cells (green, labeled with anti-GFP antisera) were prepared as per Fig. 4 with FACS for citrine, exposure to AKT1 lentivirus, replating on rat cortical cells, and cultured for 4 weeks after exposure to doxycycline (Dox) for the indicated durations. (A) Immunofluorescence for pAKT (blue) is detectable in some of the citrine+ cells after one week of exposure and increases subsequently, as quantified. (B) Immunofluorescence for mCherry in citrine+ cells parallels that of pAKT.
